# NEOage clocks - Epigenetic clocks to estimate post-menstrual and postnatal age in preterm infants

**DOI:** 10.1101/2021.05.13.444018

**Authors:** Stefan Graw, Marie Camerota, Brian S. Carter, Jennifer Helderman, Julie A. Hofheimer, Elisabeth C. McGowan, Charles R. Neal, Steven L. Pastyrnak, Lynne M. Smith, Sheri A. DellaGrotta, Lynne M. Dansereau, James F. Padbury, T. Michael O’Shea, Barry M. Lester, Carmen J. Marsit, Todd M. Everson

## Abstract

Epigenetic clocks based on DNA methylation (DNAm) can to accurately predict chronological age and are thought to capture biological aging. A variety of epigenetic clocks have been developed for different tissue types and age ranges, but none has focused on age prediction for preterm infants. Epigenetic estimators of biological age might be especially informative in epidemiologic studies of neonates, particularly those born preterm, since this is a key developmental window. Neonatal DNAm is dynamic and preterm infants are at heightened risk of developmental impairments. We aimed to fill this gap by developing epigenetic clocks for neonatal aging in preterm infants.

As part of the Neonatal Neurobehavior and Outcomes in Very Preterm Infants (NOVI) study, buccal cells were collected at NICU discharge to profile DNAm levels in 542 very preterm infants. We applied elastic net regression to identify four epigenetic clocks (NEOage) predictive of post-menstrual and postnatal age, compatible with the Illumina EPIC and 450K arrays. We observed high correlations between predicted and reported ages (0.93 – 0.94) with root mean squared errors (1.28 - 1.63 weeks).

Epigenetic estimators of neonatal aging in preterm infants can be useful tools to evaluate biological maturity and associations with neonatal and long-term morbidities.

## 1. Introduction

DNA methylation (DNAm) is one of the most studied epigenetic mechanisms and acts at the interface between the environment and human health. Changes in DNAm are also strongly correlated with aging [1] and are most dynamic during pediatric age [2]. Aging-related fluctuations in DNAm levels have been capitalized on by researchers to develop “epigenetic clocks”, sets of CpG sites whose methylation extents have been shown to accurately predict chronological age and are thought to capture biological aging [2, 3]. These predicted ages are often referred to as epigenetic age or DNAm age. Greater DNAm age relative to chronological age, also known as age acceleration (AA), has been shown to be associated with age-related phenotypes in adults, such as frailty, chronic diseases and mortality [4].

A variety of epigenetic clocks have been developed to predict numerous age metrics in different tissue types and age ranges [5]. One of the most widely used pan-tissue clocks to estimate chronological age was created by Horvath and is based on over 8,000 samples from 51 healthy tissues (age range: 0-101 years) [6]. However, DNAm age estimates from Horvath’s epigenetic clock become more precise as chronological age increases and are most variable in pediatric samples [7]. Hannum et al. developed a clock based on blood with an age range of 19-101 years [8] while other clocks are designed to capture physiological measures of biological age rather than chronological age. These include DNAm PhenoAge [9] and DNAm GrimAge [10] and are both blood-based. Many studies have successfully generated epigenetic clocks for various tissues, age ranges, and morbidities, leading to very promising predictors of chronological age in adults, and to potentially useful biomarkers for the diseases of aging. While some epigenetic clocks include children, most clocks are primarily focused on adults and extrapolating them to children results in inaccurate predictions [2, 11]. Additionally, AA metrics that are derived from these clocks may not be as relevant to the health conditions that are most important to children and adolescents. To address this issue McEwen et al. developed PedBE, an epigenetic clock that focuses on estimating chronological age of children ranging from 0 (birth) to 20 years old and is based on buccal epithelial cells [2]. However, the definition of chronological age becomes less meaningful proximal to birth and is especially skewed among infants born preterm. Infants born preterm might differ biologically from infants of the same chronologic or postnatal age that are born full-term. Epigenetic clocks, such as those developed by Knight et al. [12] or Bohlin et al. [13], have been created to capture gestational age (GA), i.e. the time from conception to birth. Both clocks are based on cord blood and therefore can only estimate gestational age, not postnatal age. To our knowledge, there exists no epigenetic clock that properly handles or is specialized for age prediction in preterm infants.

The WHO estimated 15 million infants, approximately 10% of live births, are born prematurely early every year (before completing 37 weeks of gestation) [14]. Preterm birth is not only associated with acute and long-term morbidities including chronic illnesses, brain injuries, and adverse neuromotor, cognitive, and behavioral outcomes [15], but it is also the leading cause of death worldwide among children under 5 years [14]. This leads to an immense emotional and financial burden for families and society. The Institute of Medicine reported in 2007 that the average medical costs of the first year were almost 10 times greater for preterm infants in the U.S., and results in a societal economic cost of $26.2 billion each year [16, 17].

Here, we present four NEOage (Neonatal Epigenetic Estimator of age) clocks, epigenetic clocks that are focused on age estimation of preterm infants based on their DNAm profile measured in an easily accessible tissue, buccal epithelial cells. Specifically, we investigated post-menstrual age (PMA), the time from conception to tissue collection at neonatal intensive care unit (NICU) discharge, and post-natal age (PNA, or chronological age), the time from birth to tissue collection (Figure 1). These epigenetic estimators of aging could be particularly important for preterm neonates, because they may provide insight into early life aging, reflect health and development, and provide a measure of early life risk for neonatal morbidities or long-term neurodevelopmental impairments.

**Figure 1.**
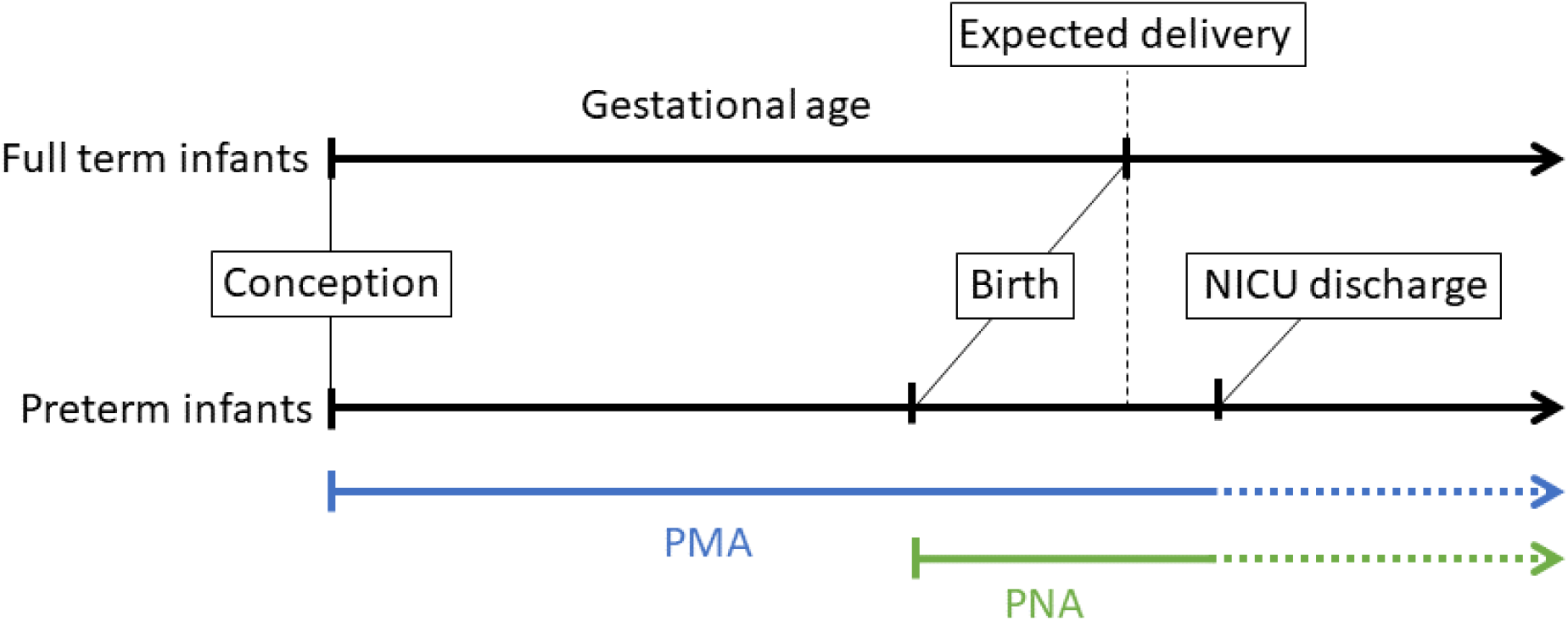
Illustration of different perinatal age metrics, measured in weeks and days, which we highlight for infants born preterm. Gestational age (GA) is defined as the time from conception to birth (expected delivery around 37-42 weeks typically refers to full-term birth, and <37 weeks refers to preterm birth). Post-menstrual age (PMA) refers to the time from conception onward, and postnatal age (PNA) is equivalent to chronological age and is the time elapsed after birth. In this study, buccal cell tissue was collected from infants at NICU discharge to profile DNA methylation.

## 2. Methods

### 2.1. Study participants

The Neonatal Neurobehavior and Outcomes in Very Preterm Infants (NOVI) study enrolled infants born <30 weeks postmenstrual age (PMA) from nine NICUs affiliated with six universities from April 2014 to June 2016. Inclusion criteria included: (1) birth <30 weeks PMA; (2) parental ability to read and speak English, Spanish, Japanese, or Chinese; and (3) residence within 3 hours of the NICU and follow-up clinic. Infants were excluded for major congenital anomalies, maternal age < 18 years, cognitive impairment, or death.

Parents of eligible infants were invited to participate in the study when an infant reached 31-32 PMA or when survival to discharge was determined to be likely by the attending neonatologist. Researchers explained study procedures and obtained informed consent in accordance with each institution’s review board. Children were included in this analysis if DNAm data was available. PMA in NOVI was calculated by adding PNA at buccal collection to the estimated gestational age at birth which was obtained via an established process [18, 19] and is described in detail by Everson et al. [15].

### 2.2. DNAm Collection and Pre-Processing

Buccal cell tissue was collected from infants that were born very preterm, at NICU discharge (Figure 1), and DNAm levels were profiled using the Infinium MethylationEPIC BeadChip (EPIC).

Genomic DNA was extracted from buccal swab samples, collected near term-equivalent age, using the Isohelix Buccal Swab system (Boca Scientific), quantified using the Quibit Fluorometer (Thermo Fisher, Waltham, MA, USA) and aliquoted into a standardized concentration for subsequent analyses. DNA samples were plated randomly across 96-well plates and provided to the Emory University Integrated Genomics Core for bisulfite modification using the EZ DNA Methylation Kit (Zymo Research, Irvine, CA), and subsequent assessment of genome-wide DNAm using the Illumina MethylationEPIC Beadarray (Illumina, San Diego, CA) following standardized methods based on the manufacturer’s protocol. The pre-processing of the data followed a modified workflow described by Everson et al. [15]. Array data were normalized via Noob normalization [20, 21] and samples with more than 5% of probes yielding detection p-values□>□1.0E-5 or mismatch between reported and predicted sex were excluded. In addition, one of two duplicated samples was omitted (we retained the duplicate sample with smallest detection p-values). Probes with median detection p-values□<□0.05, probes measured on the X or Y chromosome, probes that had single nucleotide polymorphisms (SNP) within the binding region or that could cross-hybridize to other regions of the genome were excluded [22]. Then, array data were standardized across Type-I and Type-II probe designs with beta-mixture quantile normalization [23, 24]. After exclusions, 706,323 probes were available from 542 samples for this study. These data are accessible through NCBI Gene Expression Omnibus (GEO) via accession series GSE128821.

### 2.3. Development of the epigenetic clocks

Since data from the EPIC and Infinium HumanMethylation450 BeadChip (450k) arrays are widely used in ongoing research projects, we considered two sets of data for all analyses: (1) a complete data set (706,323 probes) with logit transformed beta-values (m-values) that is compatible with EPIC arrays (hereafter referred to as the EPIC data set) and (2) a subset of the logit-transformed data (364,410 probes) that is compatible with both EPIC and 450k arrays (hereafter referred to as the 450k data set). Penalized regression models (“glmnet” function in glmnet R package [25]) were fit to both data sets to identify sets of CpGs (NEOage clocks) predictive of PMA and PNA (4 total clocks: PMA-EPIC, PNA- EPIC, PMA-450k and PNA-450k). The alpha parameter of glmnet was set to 0.5 (elastic net regression) and lambda (PMA- EPIC: 0.049, PNA-EPIC: 0.0677, PMA-450k: 0.097 and PNA-450k: 0.2038) was chosen such that the mean cross-validated error is minimized with 10-fold cross validation (“lambda.min” result from “cv.glmnet”function in glmnet R package [25]).

We fit a series of penalized regression models to both data sets (EPIC and 450k) applying leave-one-out (LOO) cross-validation. This procedure allowed us to assess prediction performances but also limit overfitting and selection bias. In LOO cross-validation, a model is trained on all but one sample to make a prediction for that held-out sample. This step is repeated until each sample is held out and predicted once and results in N potentially unique sets of CpGs for a given outcome, where N is the sample size. Because our sample contained multiple births (e.g., twins), we additionally removed all siblings from the training set of all non-singleton children. The performance of predicted age outcomes was evaluated by examining their correlation with the measured outcome and root mean squared error (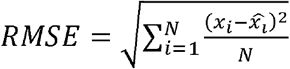 with *x_i_* and 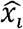 being the observed and estimated age, respectively).

In addition, prediction performances of models trained using the complete (not LOO approach) 450k data set were evaluated in an independent external data set (GSE72120 [26]) that measured DNAm in 34 preterm and 36 full-term infants using the 450k array. This data set was chosen because to our knowledge it is the closest comparable data, but it is important to point out the difference between both data sets, as one measured DNAm of buccal swabs via the EPIC array and the other profiled DNAm in saliva using the 450k array. We evaluated the performance of our PMA-450k and PNA-450k NEOage clocks in the test sample by examining the correlation between predicted and measured outcomes. We also report the RMSE.

### 2.4. Application of existing epigenetic clocks

To compare our newly-developed NEOage clocks to existing clocks, we applied Horvath’s skinblood clock [27] and PedBE [2] to estimate PNA in our data and in the independent external data set. Horvath’s skin-blood clock includes 391 CpGs and was developed with DNA from human fibroblasts, keratinocytes, buccal cells, endothelial cells, blood, and saliva (age range: 0-92 years). Out of the 391 CpGs, 345 CpGs were available in the NOVI and saliva data set. For the NOVI data set, 42 out of the 46 missing CpGs were substituted with closest CpGs within 5,000bp. The remaining 4 missing CpGs were omitted; 3 CpGs did not have CpGs available in our data that were within 5,000bp and 1 CpG was located on chromosome X (excluded during data preprocessing). Analogously for the saliva data set, 40 of the 46 missing CpGs were substituted with closest CpGs within 5,000bp. The remaining 6 missing CpGs were omitted; 5 CpGs did not have CpGs available in the saliva data set that were within 5,000bp and 1 CpG was located on chromosome X. The PedBE clock (age range: 0-20 years), developed with pediatric buccal epithelial cells, consists of 94 CpGs. There were 5 CpGs not available in the NOVI and saliva data set, which were substituted by the closest CpGs within 5,000bp. No CpGs were omitted. Performance of predicted PNA was evaluated by their correlation with the measured PNA and RMSE.

### 2.5. Enrichment analysis

To gain insights into the biological functions of the genes associated with the identified CpGs included in the four NEOage clocks, we performed an enrichment analysis. We utilized the “gometh” function in missMethyl Bioconductor package [28], that performs a hypergeometric test, while taking the number of CpG sites per gene into account. For the enrichment analysis involving the CpGs of the 450k clocks, we specified the array type to be “450k” and provided a list of CpGs that were considered (364,410 probes) for the “all.cpg” argument of “gometh”. Analogously, we specified the array type to be “EPIC” for the enrichment analysis involving the CpGs of the EPIC clocks and also provided a list of CpGs that were considered (706,323 probes). We evaluated both options for databases provided by “gometh”: GO (Gene Ontology) and KEGG (Kyoto Encyclopedia of Genes and Genomes).

## 3. Results

We applied elastic net regression to identify the sets of CpGs that are predictive of PMA and PNA in a unique population of preterm neonates. We compared the prediction performances of our NEOage clocks to two existing epigenetic clocks (Horvath’s skin-blood clock and PedBE) by evaluating their performances in our NOVI (buccal cells) data set and an external saliva data set.

### 3.1. NEOage clocks

We identified four epigenetic clocks predictive of either PMA or PNA that are compatible with the EPIC array or 450k array. The number of CpGs within each clock range from 303-522 CpGs with varying degrees of overlap between the clocks (see Figure 2). CpGs for each NEOage clock with the corresponding coefficients to calculate DNAm age are provided in the Supplementary Material (Supplementary Tables 1-4, Supplementary Code 1).

**Figure 2.**
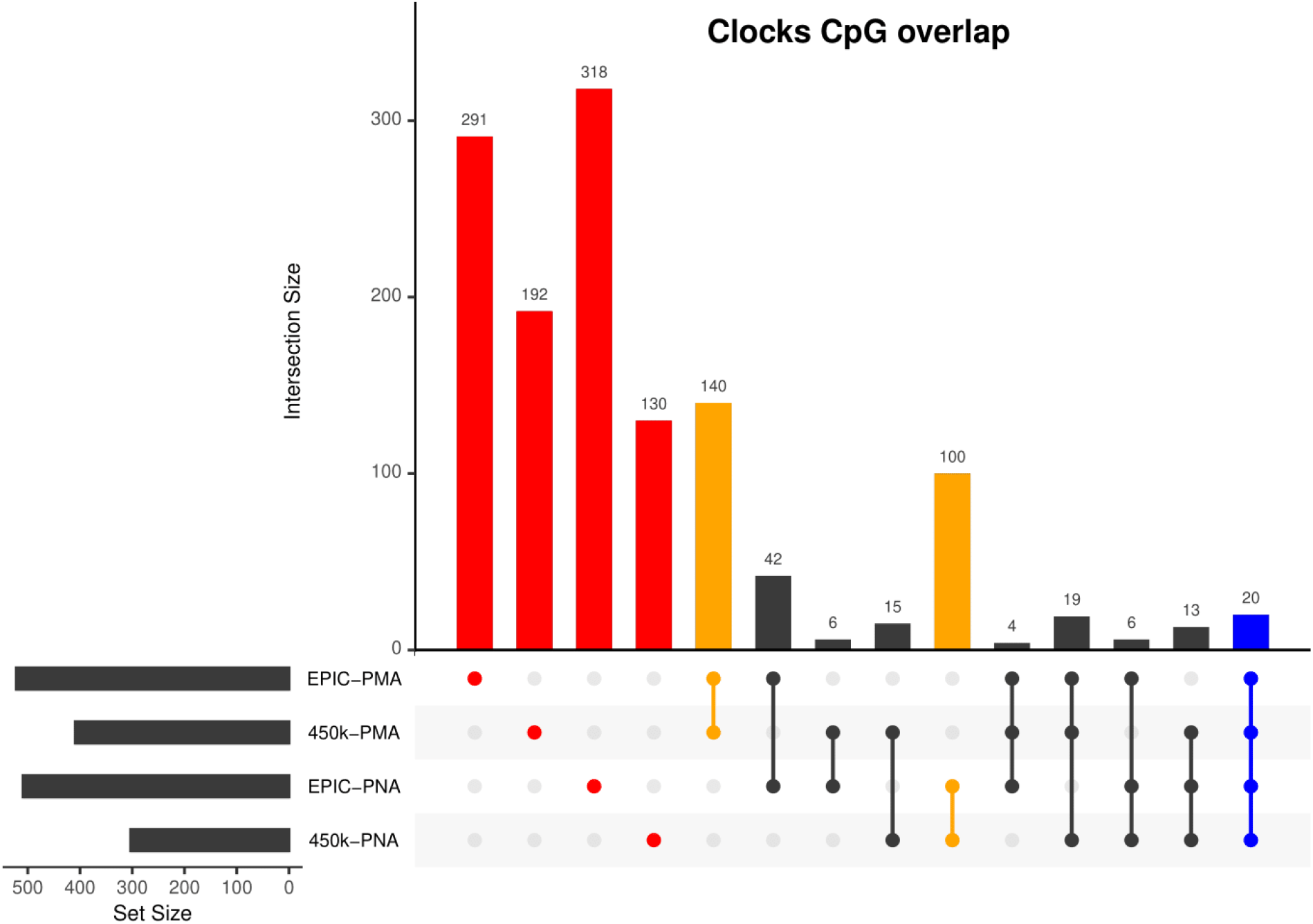
Upset plot of CpGs included in our four NEOage clocks. Highlighted in red are the number of CpGs that are unique to each individual clock. Highlighted in orange are the number of overlapping CpGs of clocks that are predictive of either PMA or PNA. Highlighted in blue are the number of CpGs that overlapped in all four clocks (additional information for the 20 common CpGs provided in Supplementary Table 5). Highlighted in black are the number of overlapping CpGs of clocks where at least one clock is predictive of PMA and at least one clock is predictive of PNA.

To assess the prediction performances without reusing information we performed LOO crossvalidation (additional information in 2.3 Development of the epigenetic clock) and evaluated prediction performances using correlations and RMSE. We observed very strong positive correlations between predicted and measured age metrics (*r* > 0.9 and p-values < 10^−16^) with very similar correlation coefficients among our four NEOage clocks (Figure 3). The predictions for PMA achieved RMSEs of 1.28 for the 450k and EPIC clocks, while predictions of PNA resulted in a RMSEs of 1.63 and 1.55, for the 450k and EPIC clocks respectively. The scatterplots in Figure 3 in combination with the strong correlations and low RMSE indicate high accuracy of our NEOage clocks.

**Figure 3.**
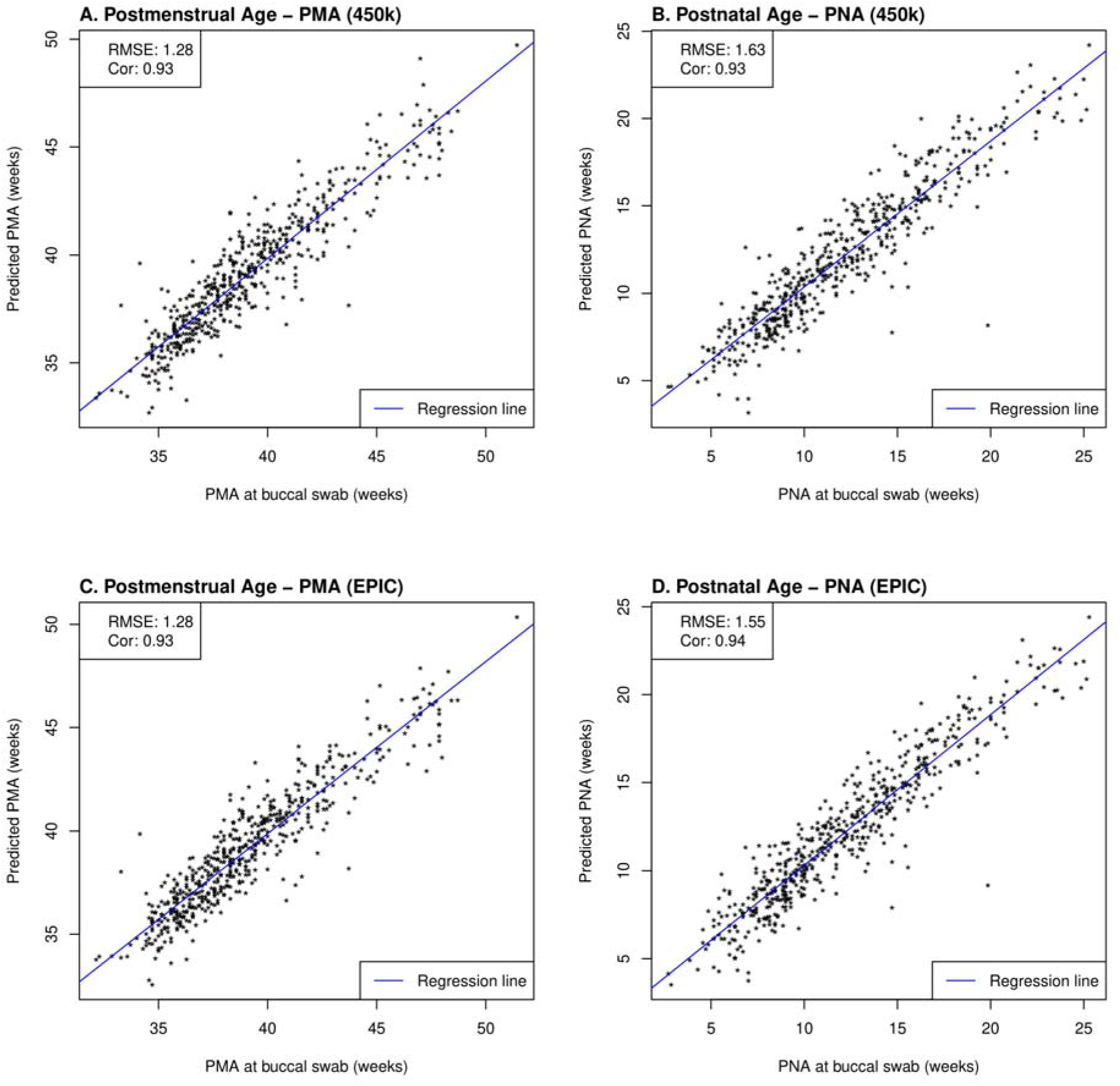
Scatterplots of estimated and measured PMA and PNA using our NEOage clocks compatible with the EPIC and 450k array within NOVI. Prediction performances are evaluated by RMSE and correlations between estimated and measured age metrics.

Next, we evaluated the prediction performance of our 450k clocks in an external independent data set that measured DNAm in saliva tissue using the 450k array. This external saliva data includes preterm and full-term infants. While Figure 4 visualizes both preterm and full-term infants, we first focused on only preterm infants in the prediction performance assessment of our NEOage clocks. Focusing on preterm infants of the saliva data allows for a more appropriate comparison of the two data sets. The prediction performances in the external saliva data set resulted in diminished but still strong correlations (PMA: r=0.61 and PNA: r=0.76), and lower RMSE for PMA (RMSE = 1.09) and similar RMSE for PNA (RMSE = 1.55), compared to the NOVI data set. However, it is important to note that the ranges of PMA and PNA in preterm infants of the saliva data are 38-42.6 and 6.9-17.6 weeks, respectively. These ranges are noticeably smaller than the ranges of PMA and PNA in the NOVI data set (PMA: 32.1-51.4 weeks; PNA: 2.7-25.3 weeks) and is likely one reason for lower correlation coefficients between predicted and reported ages in this dataset.

**Figure 4.**
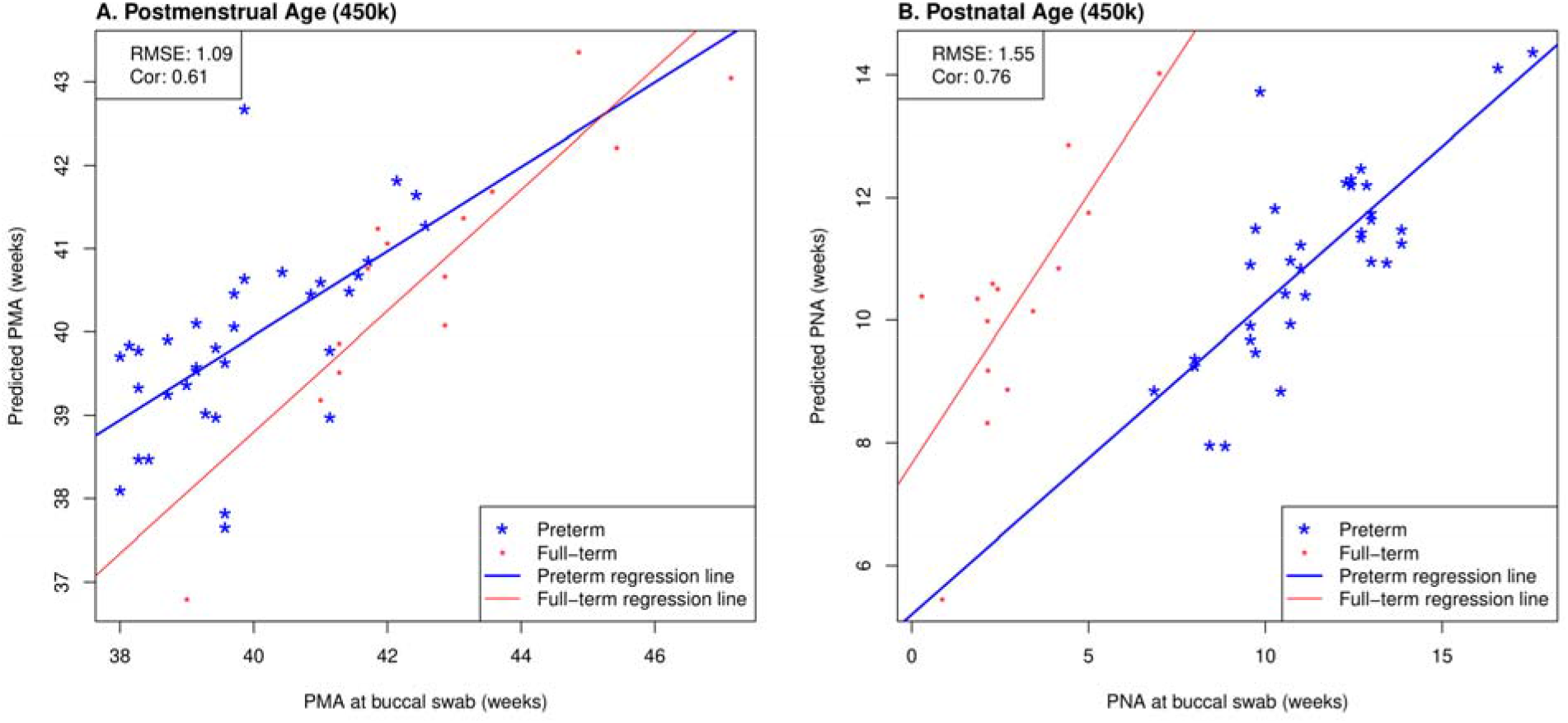
Scatterplots of estimated and measured PMA and PNA using our 450k clocks in an external saliva data set (GSE72120 [26]) that included full-term (red) and preterm (blue) infants. This saliva data set was measured by the 450k array. The reported prediction performances, RMSE and correlation coefficients, between estimated and measured age metrics are based on preterm infants only since our NOVI training data did not include any full-term infants.

While we observed strong predictive performance for our newly developed NEOage clocks, the existing Horvath skin-blood clock and PedBE clock did not predict PNA as accurately in preterm infants. As shown in Figure 5, the correlations between estimated and measured PNA are moderate in the NOVI data set (Horvath: *r* = 0.44 and PedBE: *r* = 0.59). The RMSE are greater for both clocks, with a noticeably greater RMSE for Horvath’s skin-blood clock (Horvath: *RMSE* = 49.68 and PedBE: *RMSE* = 8.68). Additionally, our NEOage clocks outperformed the existing clocks in the independent saliva data set. Analogously, Figure 6 displays both preterm and full-term infants. For preterm infants (highlighted in blue), Horvath skin-blood clock and PedBE exhibit weak correlations (Horvath: *r* = 0.31 and PedBE: *r* = 0.19) with RMSE of 38.49 and 12.93 weeks, respectively. For full-term infants (highlighted in red), the Horvath skin-blood clock correlation is *r* = 0.60 and PedBE correlation is *r* = 0.20 with RMSE of 46.31 and 5.54 weeks, respectively. Interestingly, Horvath’s clock yields a substantially better correlation between reported and predicted age for full-term infants compared to preterm infants, while the PedBE clock yielded weak correlations for both groups. Yet, while the correlations are stronger for Horvath’s clock, the actual predicted ages were closer to the reported ages for the PedBE clocks. In contrast, PNA prediction of full-term infants using our NEOage 450k PNA clock has a stronger correlation (*r* = 0.76) than both existing clocks and a similar RMSE of 7.42 weeks compared to PedBE. The best prediction performance for the full-term infants resulted from our NEOage 450k PMA clock with a correlation of 0.90 and RMSE of 2.14 weeks.

**Figure 5.**
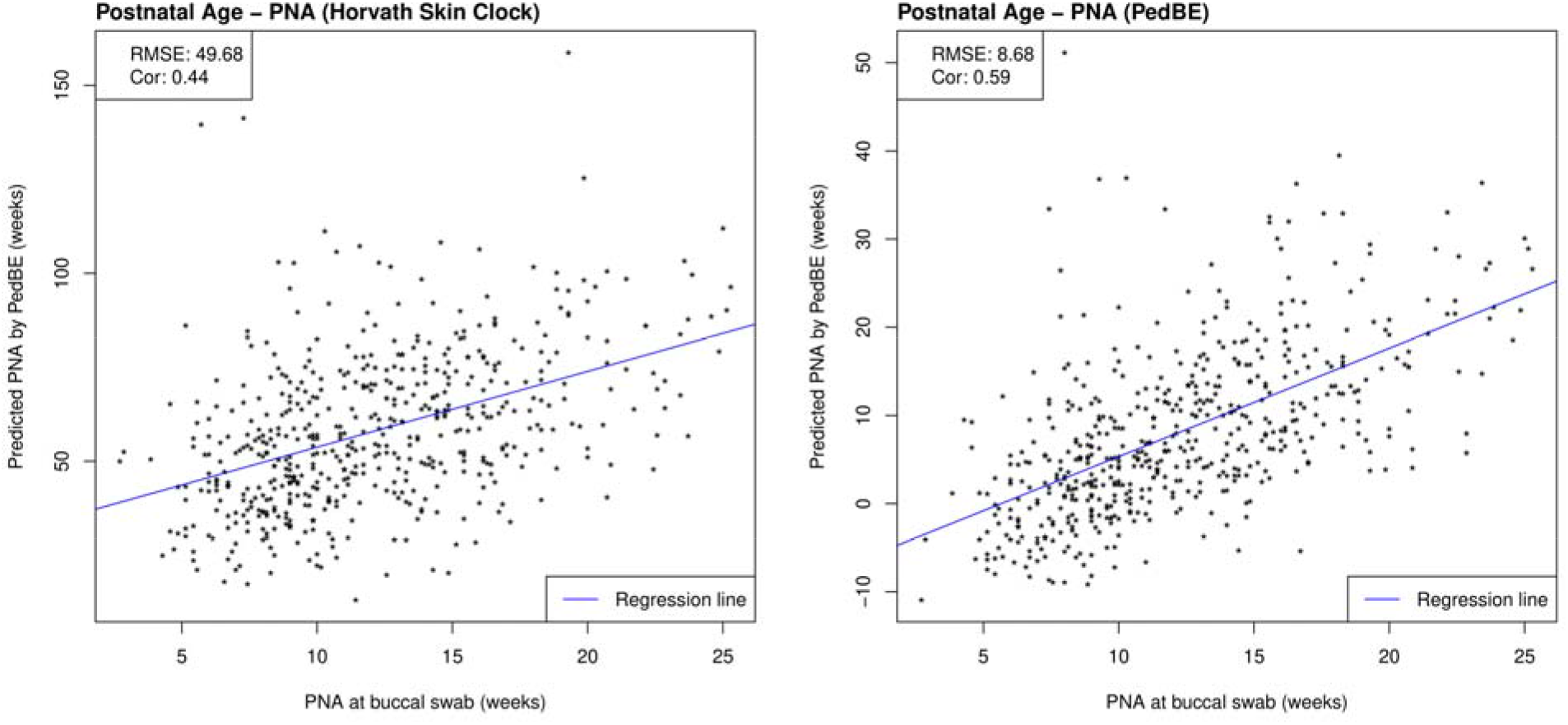
Scatterplots of PNA estimated by Horvath’s skin-blood clock and PedBE and measured PNA within NOVI. Prediction performances are evaluated by RMSE and correlations between estimated and measured PNA.

**Figure 6.**
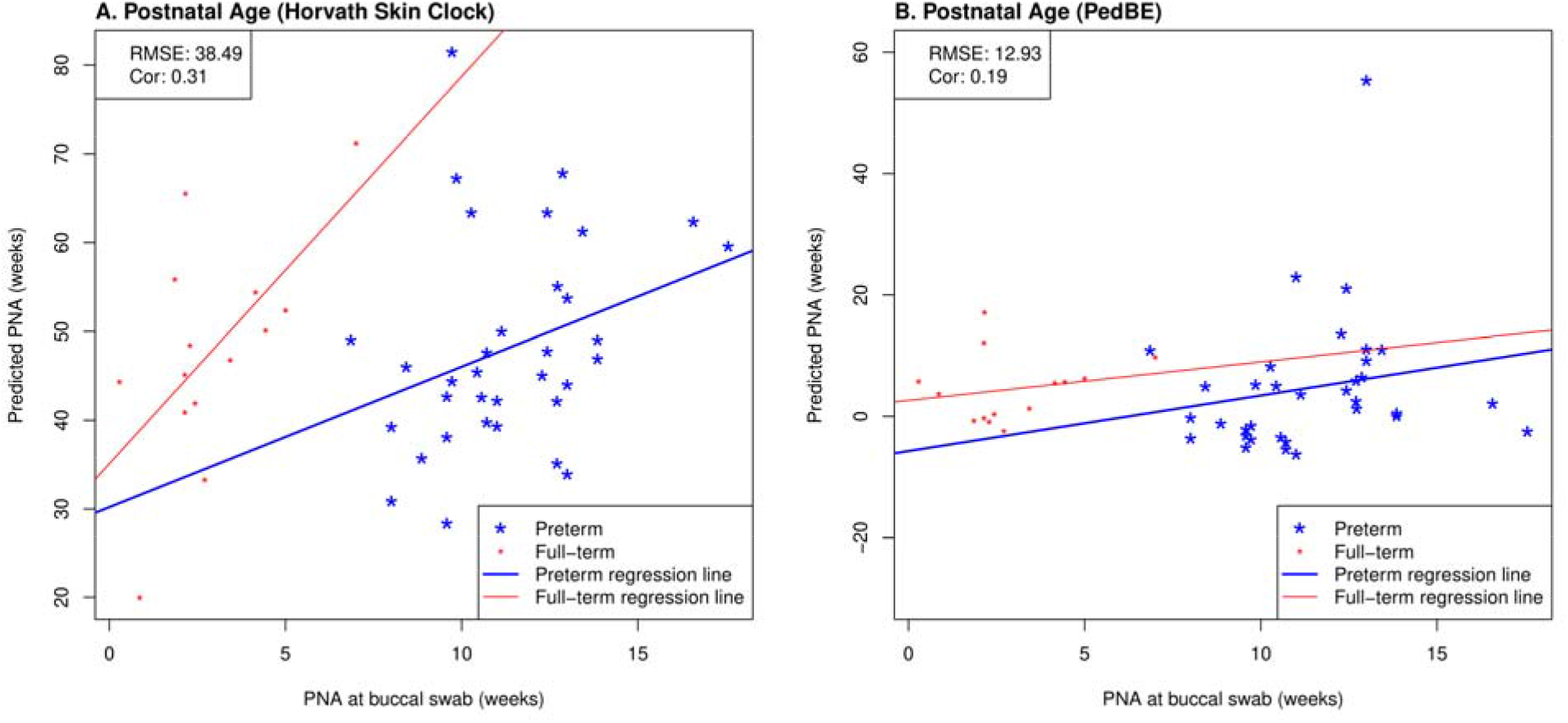
Scatterplots of measured PNA and PNA estimates by Horvath’s skin-blood clock and PedBE in an external saliva data set (GSE72120 [26]). This saliva data set was measured by the 450k array and included full-term (red) and preterm (blue) infants. The reported prediction performances, RMSE and correlation coefficients, between estimated and measured age metrics are based on preterm infants only.

### 3.2. Enrichment analysis

We performed enrichment analyses for the CpGs included in the four NEOage clocks that we characterized to evaluate potential pathway enrichments of genes associated with CpGs that we identified. No pathways or GO terms were significantly enriched after FDR correction (FDR < 0.1), but the KEGG pathways and GO terms that tended to have the smallest raw p-values included immune and inflammatory responses, endocrine activities, steroidogenesis, cellular proliferation, cellular differentiation and organization, and organ morphogenesis. Tables containing the 20 most significantly enriched pathways are provided in Supplementary Material (Supplementary Table 6-13).

## 4. Discussion

While there has been some progress in addressing the lack of epigenetic clocks focusing on pediatric populations in recent years [2], to our knowledge, there currently exists no epigenetic clock that is specialized for preterm infants, nor for age prediction specific to the neonatal period. Preterm infants present a unique population due to the shift of their biological and chronological age progress relative to full-term infants. To fill this gap, we developed four NEOage clocks that are based on preterm infants from the NOVI study to estimate PMA and PNA (EPIC- and 450k-compatible) and include 303-522 CpGs. We demonstrate that our newly developed NEOage clocks outperform two established epigenetic clocks, Horvath’s skin-blood clock and PedBE, both in our NOVI buccal data set and in an external saliva data set of infants that were born preterm.

A systematic deviation of full-term infants can be observed in Figure 4 and Figure 6. This shift appears to be more dominant in PNA predictions and might indicate that our PMA and PNA clocks capture a similar aging signature, but that our PNA clocks are more sensitive to the gestational age at birth. Pre- and full-term infants, as shown in Figure 4B, appear to have moderately similar regression slopes, but different intercepts, which is most likely a result of their different gestational age at birth. While extrapolation of our NEOage clocks outside of their training range is not recommended, it can be expected that prediction accuracy decreases with greater age differences (similar to extrapolating adult clocks to children, or pediatric clocks to the neonatal period). However, if extrapolation of age outside of our training age range but proximal to birth is necessary, our PMA clocks might be more appropriate.

We observed noticeable differences in RMSE when comparing reported ages to predicted ages from existing clocks [2, 6], predominantly in estimates from Horvath’s skin-blood clock, but also PedBE. One possible explanation is that both clocks were not specifically developed for this age range. For these existing clocks, age is estimated in years, which was then transformed to weeks by multiplying by 52. Hence, any prediction errors might be amplified. In addition, PNA is greatly overestimated for all infants by Horvath’s skin-blood clock, meaning that estimated PNA is greater than measured PNA for every infant.

While PMA seems to provide a more generalizable estimate of age, it comes with the limitation that the day of conception (reference point to calculate PMA) is not as precise of a measurement as day of birth (reference point to calculate PNA) and therefore is associated with a certain degree of uncertainty. Another limitation is the extension of these clocks to other tissue types, because our NEOage clocks are based on buccal cells collected via cheek swabs from preterm infants. Generalizing our NEOage clock to different tissue types will most likely compromise the prediction performance. Nevertheless, buccal swab is minimally invasive and thus is specifically important in pediatric and neonatal populations where more invasive sampling may deter study participation [29]. While blood samples provide large amounts of DNA with good quality, it requires an invasive and expensive procedure with technical difficulties, can be difficult or impossible to collect from preterm neonates, and causes discomfort and increased risk of infection [29]. In addition, buccal epithelial cells have been shown to be better proxy for the brain than peripheral blood [30]. The collection of buccal cells and saliva is less complicated, inexpensive and non-invasive [29], with the added benefit of buccal cells being less heterogeneous [2, 30]. A possible contamination of prenatal fetal sample with maternal cells can be avoided by performing a short terminal repeats analysis [29].

With our newly developed NEOage clocks we aim to fill the gap of methylation clocks trained on pediatric samples [31] and based on buccal cells, an easily accessible tissue that requires no invasive procedures.

Our epigenetic estimators of neonatal aging in preterm infants might be particularly valuable in this population of neonates, because it could allow us to gain insight into early life aging and reflect influences on subsequent health and development. Further, establishing precise estimators of PMA might help us to develop tools to more accurately determine the day of conception and measurements associated with it (e.g. PMA and gestational age).

## 5. Conclusion

We have introduced our four NEOage clocks that are specific to the assessment of epigenetic age in very preterm neonates. Our NEOage clocks are based on buccal cells, a tissue that is easily accessible and requires no invasive intervention. Postmenstrual age (PMA) and post-natal age (PNA) can be accurately estimated utilizing DNAm measured by either the Illumina 450k or EPIC array. We demonstrated that our NEOage clocks outperform two existing clocks by assessing their prediction performances in two preterm infant data sets. With our NEOage clocks, we have provided tools to examine neonatal aging, age acceleration and their association with neonatal health and development in a unique population of very preterm infants.

## Data Availability Statement

The DNA methylation data generated in the current study are available in the NCBI GEO via accession series GSE128821. R codes used for the analyses presented in the paper are available upon request to the corresponding author.

## Supplementary Material

**Supplementary Table 1** CpGs and corresponding coefficients of the 450k-PMA NEOage clock

**Supplementary Table 2** CpGs and corresponding coefficients of the 450k-PNA NEOage clock

**Supplementary Table 3** CpGs and corresponding coefficients of the EPIC-PMA NEOage clock

**Supplementary Table 4** CpGs and corresponding coefficients of the EPIC-PNA NEOage clock

**Supplementary Table 5** Annotation of the 20 common CpGs of the NEOage clocks

**Supplementary Table 6** Top 20 pathways from GO pathway analysis for CpGs included in the 450k-PMA NEOage clock

**Supplementary Table 7** Top 20 pathways from KEGG pathway analysis for CpGs included in the 450k-PMA NEOage clock

**Supplementary Table 8** Top 20 pathways from GO pathway analysis for CpGs included in the 450k-PNA NEOage clock

**Supplementary Table 9** Top 20 pathways from KEGG pathway analysis for CpGs included in the 450k-PNA NEOage clock

**Supplementary Table 10** Top 20 pathways from GO pathway analysis for CpGs included in the EPIC-PMA NEOage clock

**Supplementary Table 11** Top 20 pathways from KEGG pathway analysis for CpGs included in the EPIC-PMA NEOage clock

**Supplementary Table 12** Top 20 pathways from GO pathway analysis for CpGs included in the EPIC-PNA NEOage clock

**Supplementary Table 13** Top 20 pathways from KEGG pathway analysis for CpGs included in the EPIC-PNA NEOage clock

**Supplementary Code 1** R code example to calculate DNAm age using NEOage clocks

