## Supplementary Code 1 for "NEOage clocks - Epigenetic clocks to estimate post-menstrual and postnatal age in preterm infants": Supplementary Code 1 NEOage.R

```
library(minfi) # logit2

# logit transformed DNA methylation data (datM; columns: samples, rows: CpGs)
datM = logit2(datM)


# PMA EPIC
## reading in clock CpGs with coefficients
NEOage_EPIC_PMA_clock=read.csv("05-13-2021_PMA-EPIC.csv", header = T) 

## subset and transform DNAm data to selected CpGs
dataClock = data.frame(t(datM[NEOage_EPIC_PMA_clock$ID[-1],]))

## calculate NEOage 
NEOage_EPIC_PMA = as.numeric(NEOage_EPIC_PMA_clock[1,2] + as.numeric(as.matrix(dataClock) %*% as.numeric(NEOage_EPIC_PMA_clock$coef[-1])))


# PMA 450k
## reading in clock CpGs with coefficients
NEOage_450k_PMA_clock=read.csv("05-13-2021_PMA-450k.csv", header = T) 

## subset and transform DNAm data to selected CpGs
dataClock = data.frame(t(datM[NEOage_450k_PMA_clock$ID[-1],]))

## calculate NEOage 
NEOage_450k_PMA = as.numeric(NEOage_450k_PMA_clock[1,2] + as.numeric(as.matrix(dataClock) %*% as.numeric(NEOage_450k_PMA_clock$coef[-1])))


# PNA EPIC
## reading in clock CpGs with coefficients
NEOage_EPIC_PNA_clock=read.csv("05-13-2021_PNA-EPIC.csv", header = T) 

## subset and transform DNAm data to selected CpGs
dataClock = data.frame(t(datM[NEOage_EPIC_PNA_clock$ID[-1],]))

## calculate NEOage 
NEOage_EPIC_PNA = as.numeric(NEOage_EPIC_PNA_clock[1,2] + as.numeric(as.matrix(dataClock) %*% as.numeric(NEOage_EPIC_PNA_clock$coef[-1])))


# PNA 450k
## reading in clock CpGs with coefficients
NEOage_450k_PNA_clock=read.csv("05-13-2021_PNA-450k.csv", header = T) 

## subset and transform DNAm data to selected CpGs
dataClock = data.frame(t(datM[NEOage_450k_PNA_clock$ID[-1],]))

## calculate NEOage 
NEOage_450k_PNA = as.numeric(NEOage_450k_PNA_clock[1,2] + as.numeric(as.matrix(dataClock) %*% as.numeric(NEOage_450k_PNA_clock$coef[-1])))
```
